# Reading the canid skeletal story: Coxofemoral joint pathology, and suggested implications for the phylogenetic and natural history of taxa

**DOI:** 10.1101/2022.09.20.508586

**Authors:** Dennis Lawler, Basil Tangredi, Christopher Widga, Michael Etnier, Terrance Martin, Luci Kohn

## Abstract

We evaluated subtle-to-incipient pathology traits in coxofemoral joints from dry bone museum specimens of: *Vulpes lagopus; Vulpes; Nyctereutes procyonoides; Urocyon cinereoargenteus; Canis lupus familiaris*; and *Canis latrans*. Multiple intra-articular structures were evaluated on acetabula and proximal femora. Primary observations included multifocal, variable osteophyte-like formations; osteophyte-like rimming of articular margins and femoral head (*ligamentum teres* attachment); and rough or worn bone. Within limitations on valid statistical applications, we observed little difference among the high trait frequencies across taxa, aligning with previous morphological observations.

Additionally, for this study, we evaluated the known history of the taxa, from deep time to the present, to consider our data in a phylogenetic context. Potential introgression over the evolution of Canidae, along with early history of the canid genome, likely supported broad and deep conservation of pathophysiological processes associated with observable pathology at the same intra-articular foci, across taxa. We also evaluated the “modern” natural histories of the taxa, noting that coxofemoral joint impacts of their respective life habits did not appear to influence pathology trait outcomes differentially.

We conclude that conservation of the physiology underlying subtle and incipient coxofemoral joint pathology that did not segregate among taxa. We hypothesize that the intersecting basic biology of growth-development and insult response, over long geological time, may owe in part to the evidently long histories of hybridization and generally high historical gene flow, with high levels of heterogeneity.

These data argue for new research to advance an interdisciplinary, integrated understanding of relationships among canid growth-development, incipient-to-subtle joint pathology, influences of natural histories across related taxa, and implications for genomic interrelationships.

## INTRODUCTION

<u>Family</u> Canidae is thought to have diverged from stem carnivorans earlier taxa c. 40 Mya, during the late Eocene (Tedford et al., 2009). Of the three <u>subfamilies</u> within Canidae, Hesperocyoninae appeared first, peaked in diversity over c. 32 – 21 Mya, and became extinct by c. 15 Mya. Subfamilies Borophaginae and Caninae appear to have derived from an early divergence with Hesperocyoninae, followed by another divergence that further separated Borophaginae from Caninae. The subfamily Borophaginae were extant over c. 34 – 2 Mya, while subfamily Caninae appears to have evolved more slowly, beginning c. 34 Mya but flourishing only after c. 10 – 8 Mya, into the present (See Tedford et al., 2009, Fig. 2).

**Figure 1.**
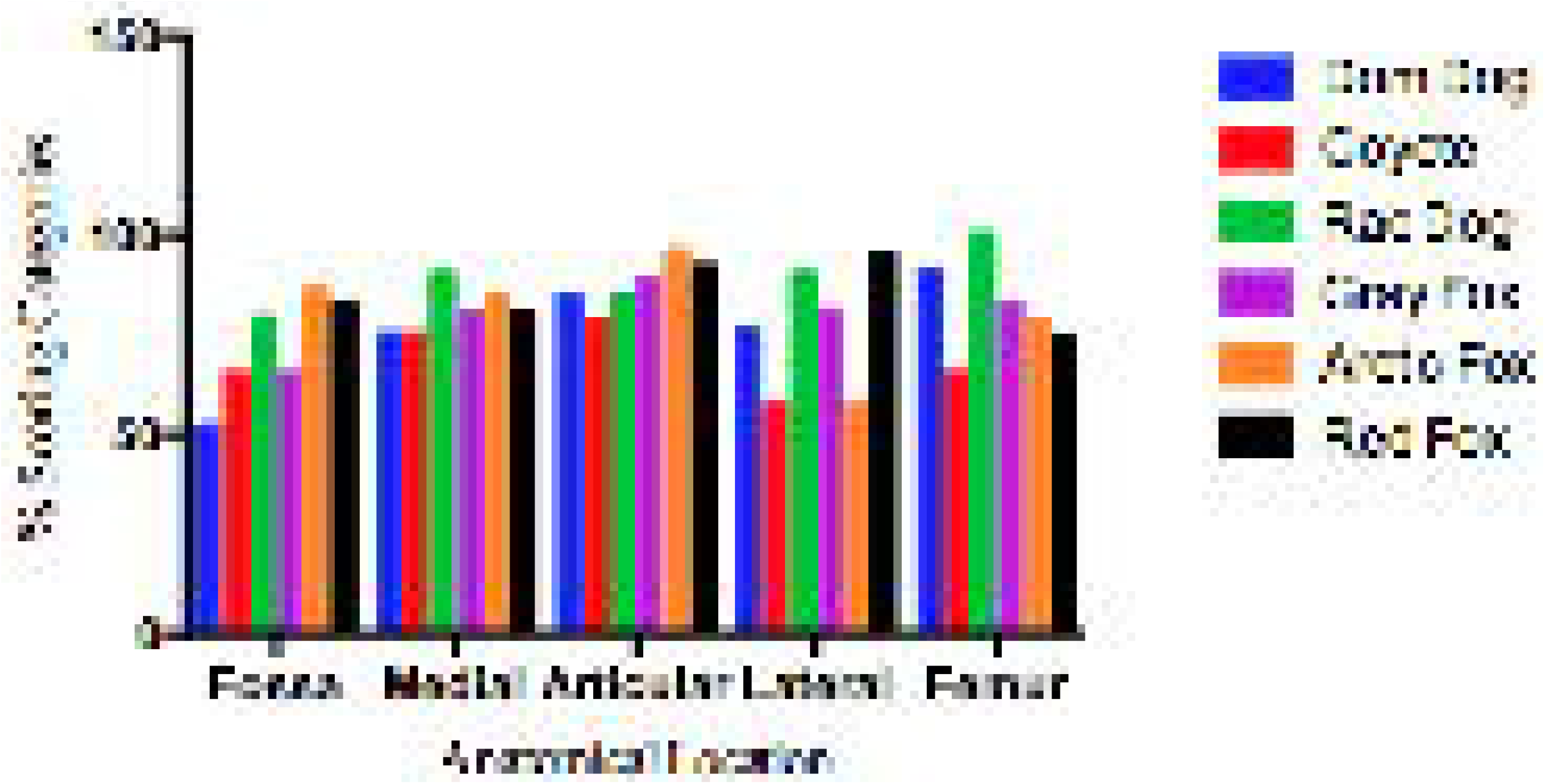
Scoring for Acetabular Fossa. **Left** or **Right** **A)** Ligament attachment

a. boney surface only
b. remnant enthesophytes round ligament attachment
c. full fossa surface osteophytes, uniform **B)** Accessory ligament groove

a. osteophytes present
b. osteophytes absent **C)** Non-ligament fossa surface

a. boney surface only
b. light osteophyte cover
  i. cranial
  ii. dorsal
  iii. caudal
  iv. ventral (some ventral and central = enthesophyte)
  v. central (some ventral and central = enthesophyte)
c. heavy osteophyte cover
  i. cranial
  ii. dorsal
  iii. caudal
  iv. ventral
  v. central **D)** Fossa shape

a. unremarkable
b. wide
  i. full
  ii. dorsal
  iii. central
  iv. ventral
c. narrow
  i. full
  ii. dorsal
  iii. central
  iv. ventral
d. skewed
  i. cranial
  ii. craniodorsal
  iii. caudal
  iv. caudodorsal

**Figure 2.**
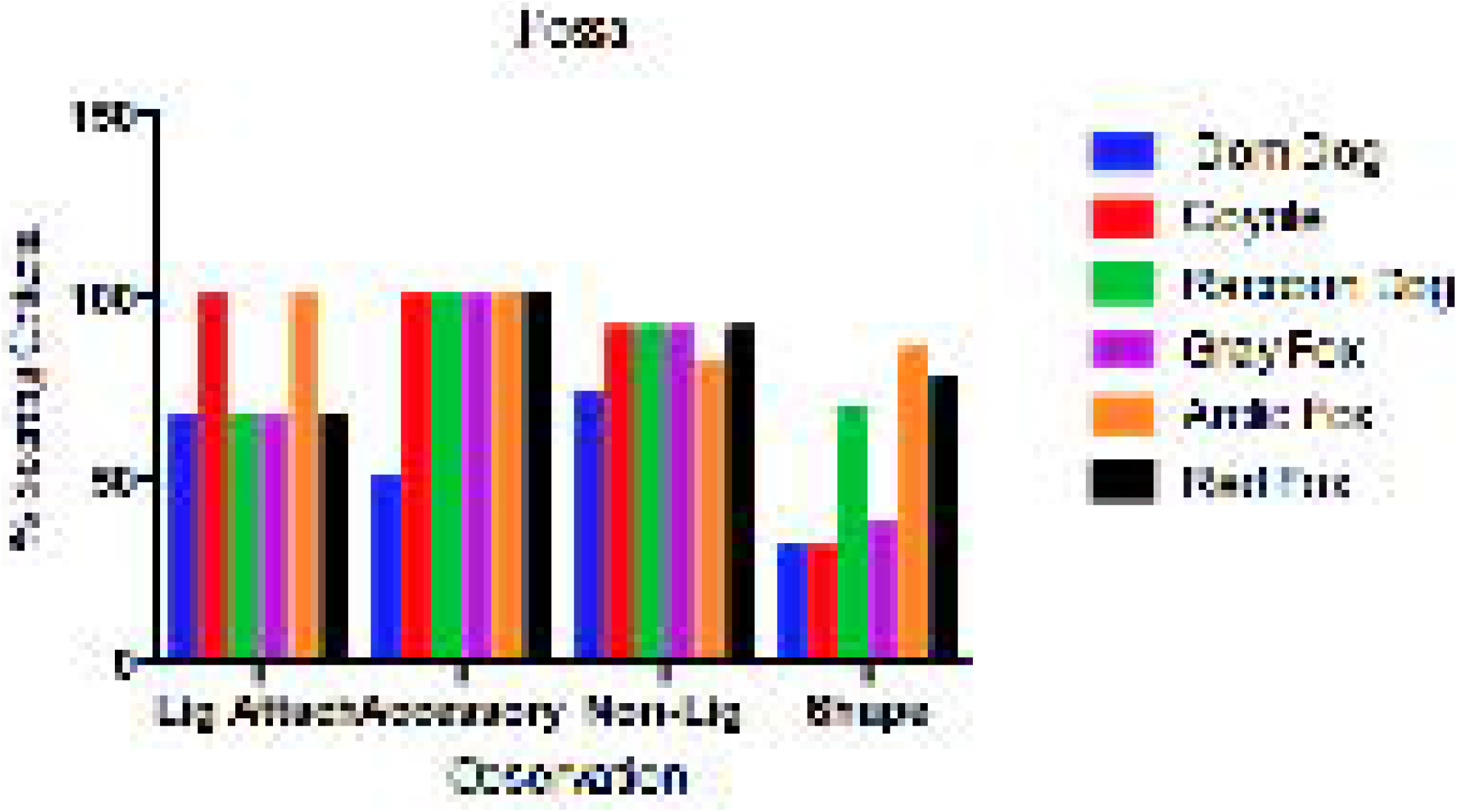
Scoring for Medial Articular Margin. **Left** or **Right** **A)** Extent of margin

a. no pathology
b. full margin
c. intermittent focused pathology at:
  i. cranial
  ii. craniodorsal
  iii. dorsal
  iv. caudodorsal
  v. caudal **B)** Character of margin

a. smooth
b. rough; irregular; variable
c. minimal
d. wide
e. impinging
  i. cranial
  ii. craniodorsal
  iii. dorsal
  iv. caudodorsal
  v. caudal
  vi. ventral
f. obliterated
g. porous

However, based on molecular data, such as those presented by Nyakatura and Bininda-Emonds (2012), the origin of <u>Family</u> Canidae was placed much earlier in time, at slightly over 60 Mya. Furthermore, using cladistic analysis, Tomiya (2011) suggested the earliest <u>caniform-feliform</u> divergence to be either c. 47 Mya or c. 38 Mya, depending on the ultimate assignment of *Miascis silvestris* (recalling that Tedford et al. (2009) suggested the origin of <u>Family</u> Canidae in that same approximate time frame.)

Some confusion is not uncommon when comparing results from multiple analytical approaches to the same data. Method-related influences in deep time can include the inevitable constraints due to: (a) Limited availability of skeletal remains (Tomiya 2011); (b) mathematical assumptions necessary for analytical processes; (c) effects of extrinsic variables that no longer can be detected (Nowak 2003); and (d) what constitutes an appropriate population sampling over deep time (Hohenlohe et al., 2017). An important corollary is that Canidae once was more varied than can be observed at present (Nowak 2003). The six taxa that are the subject of this report have complex phylogeny, characterized generally by rather high gene flow, high heterogeneity, and degrees of hybridizing (Gopalakrishnan et al., 2018).

From previous observations (Lawler et al., 2021a; Lawler et al., 2021b), we had hypothesized deep-time governance of joint biology via preserved and shared interacting physiological pathways that are involved with joint growth and development, as well as insult response (Tangredi and Lawler 2019). Our present observations suggest that “range of normal” morphology surrounding joint growth and development, and (sometimes concurrent) responses to early joint insults, should be regarded as foundationally similar (Tangredi and Lawler, 2019; Lawler et al., 2021a; Lawler et al., 2021b).

## MATERIALS AND METHODS

The population from which these specimens were selected included ancient and modern skeletal remains of 88 canids from 6 taxa. The skeletal remains are curated at: The Illinois State Museum Research and Collections Center, Springfield IL (ISM); Illinois State Archaeological Survey, Champaign IL (ISAS); East Tennessee State University Museum of Natural History, Johnson City TN (ETMNH); Michael Etnier collection, Western Washington University, Bellingham WA (MEWWU); and University of Washington Burke Museum, Seattle WA (UWBM).

### Taxa

This study included (subfamily Caninae, tribe canini; n=24): Ancient and modern domestic dog (*Canis lupus familiaris*; n=15); and ancient and modern coyote (*Canis latrans*; n=9). Vulpines (subfamily Caninae, tribe vulpini; n=64) included: Modern red fox (*Vulpes;* n=30); modern arctic fox (*Vulpes lagopus;* n=9); and modern raccoon dog (*Nyctereutes procyonoides*, n=16). Gray foxes (*Urocyon cinereoargenteus;* n=9) also were included. *Urocyon* is the most basilar among vulpini that remain extant. The history is unique, extending deeper into geological time (Goddard et al., 2015). While not all of the dating and forward progression of *Urocyon* from its origin are agreed upon universally, for this report, we analyzed *Urocyon cinereoarenteus* with the vulpini.

### Procedures

Specimen selection from a large database (DFL) was based on ability to describe and score pathology features from earlier photographs (Canon SX720 HS, Canon Inc., Melville NY 11747). Acetabular structures of interest included fossa, medial and lateral articular margins, articular surface, and peri-articular bone, with selection for this study depending particularly on visualizing the *ligamentum teres* proximal and distal attachments. Acetabular structures that were evaluated include fossa, medial articular margin, articular surface, lateral articular margin, and periarticular bone. Proximal femoral structures that were evaluated included articular surface and margin, and peri-articular features. This approach was chosen to allow evaluation of ligaments-pecific pathology associations in the acetabular fossa and on the femoral head, thus ensuring that all relevant parts of the specimen would be visible from the photographs.

### Evaluation and Data Analyses

We described observed pathology by location, gross appearance, and spatial relationships (Mustonen et al., 2017; Lawler et al., 2021a; Lawler et al., 2021b), according to a predetermined protocol (Figs. 1–5). Where necessary, magnification or backlighting were used to support examination. The features of interest were scored individually as present or absent, and irrespective of sex, chronological age, and severity. Sex of specimens frequently was not available, and mutual confounding can occur between chronological age and severity. Since we worked with cleared bone from museum skeletal remains, we did not require imaging beyond photography. No live animals were used, and no animals were euthanized for this study.

**Figure 3.**
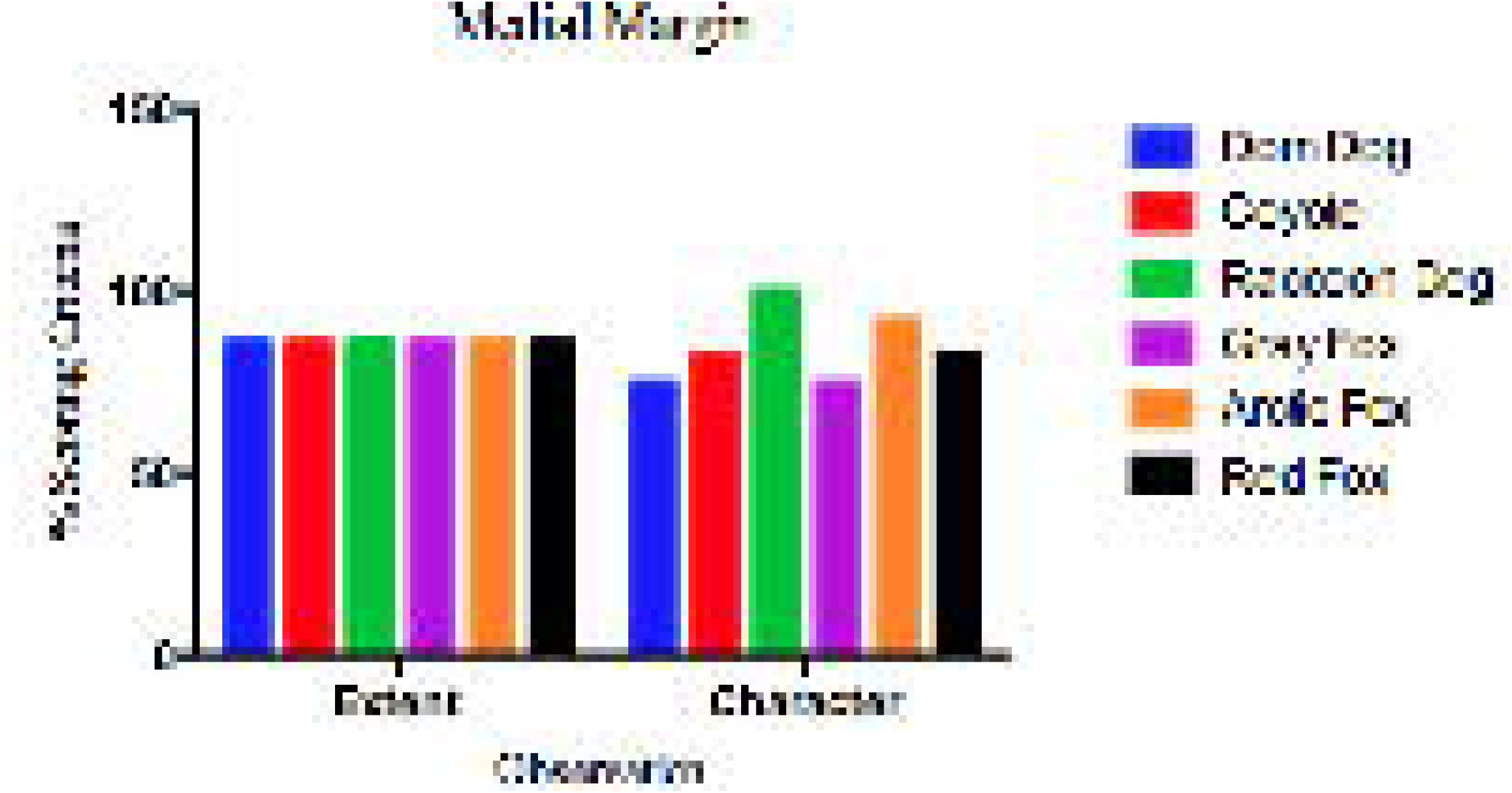
Scoring for Articular Surface. **Left** or **Right** **A)** Surface character

a. smooth
b. rough
c. depression
d. wear
e. wide
f. osteophytes flat
g. osteophytes raised
h. bone loss
i. linear
j. porous
k. irregular
l. striated
m. eburnation **B)** Spatial medial-lateral

a. medial
b. central
c. lateral **C)** Spatial cranial–caudal

a. cranial
b. craniodorsal
c. dorsal
d. caudodorsal
e. caudal
f. ventral **D)** Distribution

a. focal
b. multifocal
c. focally disseminated
d. coalescing
e. disseminated

**Figure 4.**
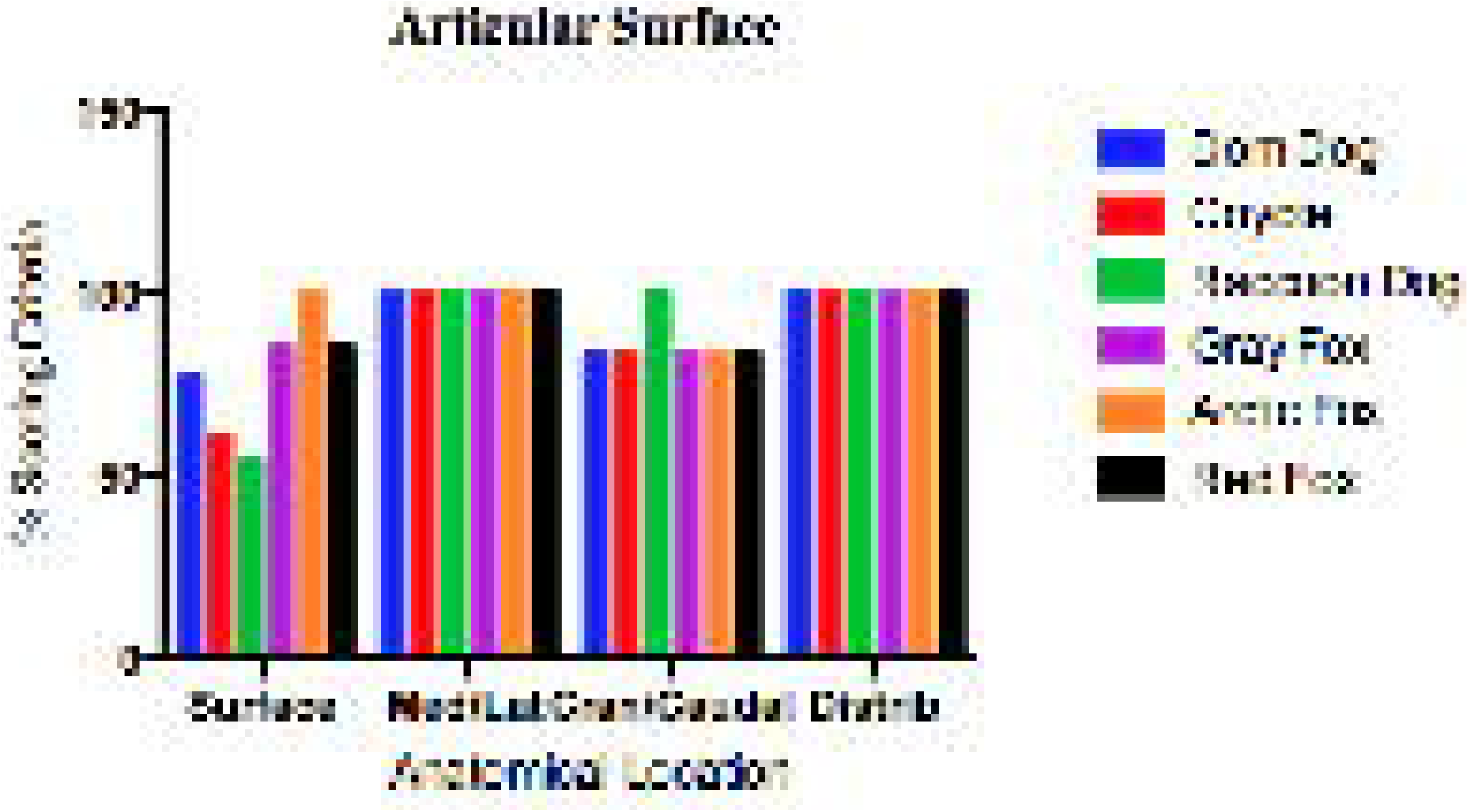
Scoring for Lateral Articular Margin. **Left** or **Right** **A)** Extent of margin

a. full margin
b. intermittent focused pathology toward cranial-dorsal
c. intermittent focused pathology toward caudal-dorsal
d. variable margin **B)** Character of margin

a. smooth
b. rough; irregular; variable
c. minimal
d. wide, thick
e. impinging-obliteration:
  i. cranial
  ii. craniodorsal
  iii. dorsal
  iv. caudodorsal
  v. caudal
  vi. ventral
f. rim
g. sharp **C)** Periarticular osteophytes

a. none
b. osteophytes present
  i. cranial
  ii. dorsal
  iii. caudal
  iv. ventral

**Table 5.**
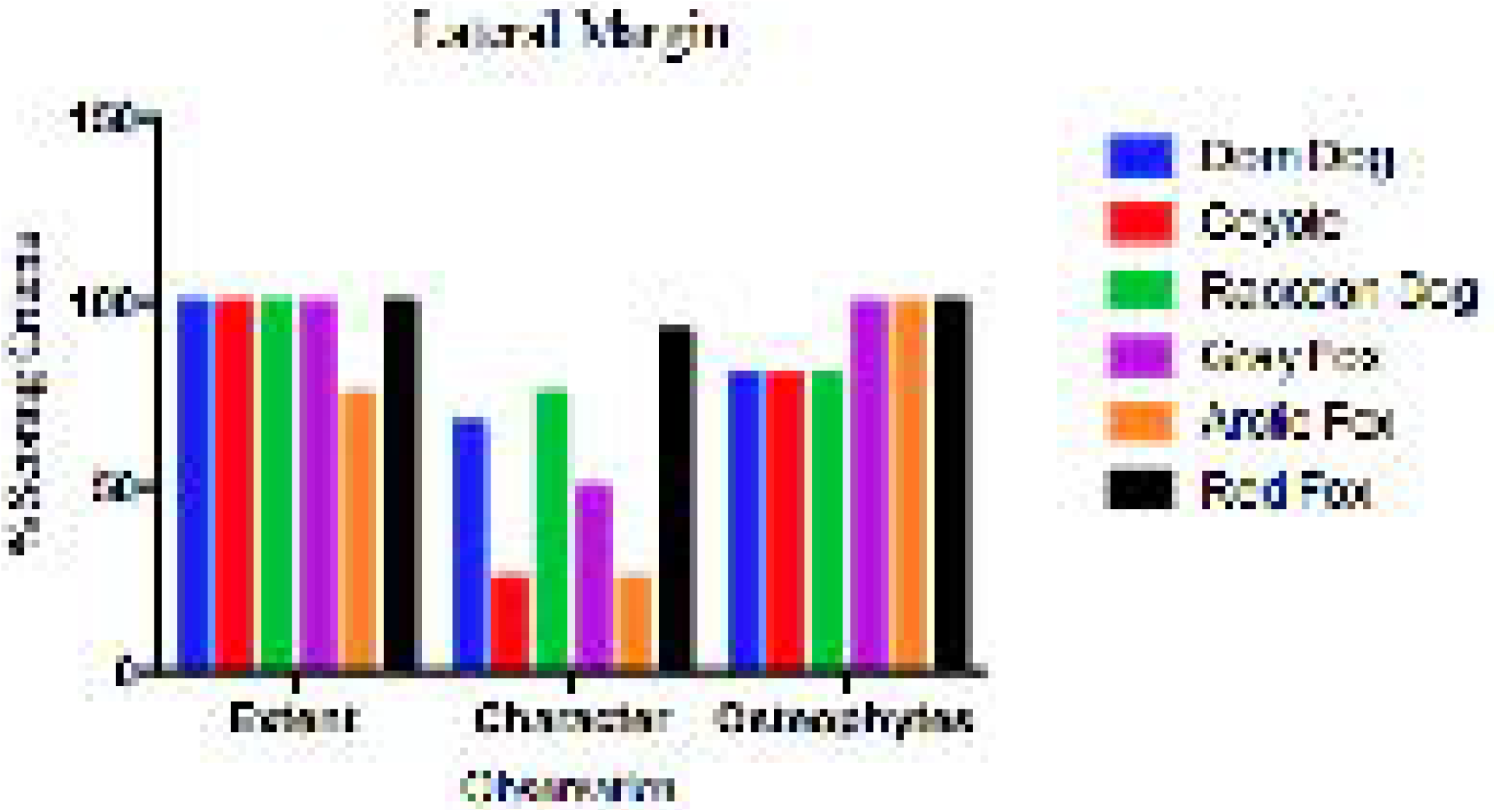
Scoring for Proximal Femur. **Left or Right** **A)** Fovea capitis

a. small size
b. large size
c. irregular shape
d. shallow
e. margin
  i. normal
  ii. rim full
  iii. rim intermittent, irregular
  iv. rim low
  v. rim severe
f. smooth floor and/or walls
g. floor - walls rough and/or osteophytes
h. bone loss
i. ossified
j. deep
k. raised **B)** Distal non-articulating component

a. smooth
b. rough; irregular; bone loss
c. depression or shallowing
d. contiguous with fovea
e. closed
f. rimmed or prominent
g. osteophytes floor or margin(s)
h. raised

The database was heterogeneous with regard to numbers within taxa and geological age. The data were mixed single and bilateral specimens. The nature of the database necessarily limited statistical comparisons. Thus, we elected to calculate the percentage of the total number of scoring categories that were represented in the database for each taxon, based on the five predetermined scoring protocols (Figs. 1–5; Supplementary Table 1). The data were evaluated from each of the category and subcategory percentages.

<u>As an illustrative example</u>, the *acetabular fossa* of the domestic dog revealed 29 possible pathological features, summarily, over subcategories A-D (Fig. 1; Supplementary Table 1). Dog specimens revealed pathology features in 15 of those 29 categories, overall (52%). For subcategory A – lesion(s) were observed in 2 of 3 possibilities (yielding a score of 67%). For subcategory B – lesion(s) were observed in 1 of 2 possibilities (yielding a score of 50%). For subcategory C – lesion(s) were observed in 8 of 11 possibilities (yielding a score of 73%). For subcategory D – lesion(s) were observed in 4 of 13 possibilities (yielding a score of 31%). This pattern of analysis was used for all five scoring protocols (Figs. 1–5), over all six taxa. Within these mathematical constraints, we considered how each taxon displayed the defined range of possible pathology. The data were not lateralized, since we have shown previously (Lawler et al., 2021b) that lateralizing analysis did not yield right-left differences.

## RESULTS

Most data were available bilaterally. All taxa expressed a majority of the possible range of pathological features, for each anatomical structure of interest. Given the wide scope of “present” observations across taxa and scoring categories, we were unable to define unique involvements for the reference point *ligamentum teres* attachments on the acetabular fossa or femoral head.

The high prevalence of pathology features across Canidae is clear from a graphical summary (Fig. 6). In some instances, focally disseminated-to-coalescing pathology partly or fully obscured other observations. In Fig. 6, a high overall presence can be seen for observations across the acetabular fossa, medial and lateral articular margins, article surface, and proximal femur, with numerical estimates ranging over 50-100% of possible outcomes. Visual obstruction of one or more features by another feature resulted in some negative outcomes for the obscured structure(s). Therefore, across the database, all interpretations must be made in the light of high overall prevalence of target observations, with some lower scores consequent to visual obstruction by other features. For example, the domestic dog seemed to have lower frequency of acetabular fossa pathology, but the above-described scoring caveats should be applied (Fig. 6, 7; Supplementary Table 1).

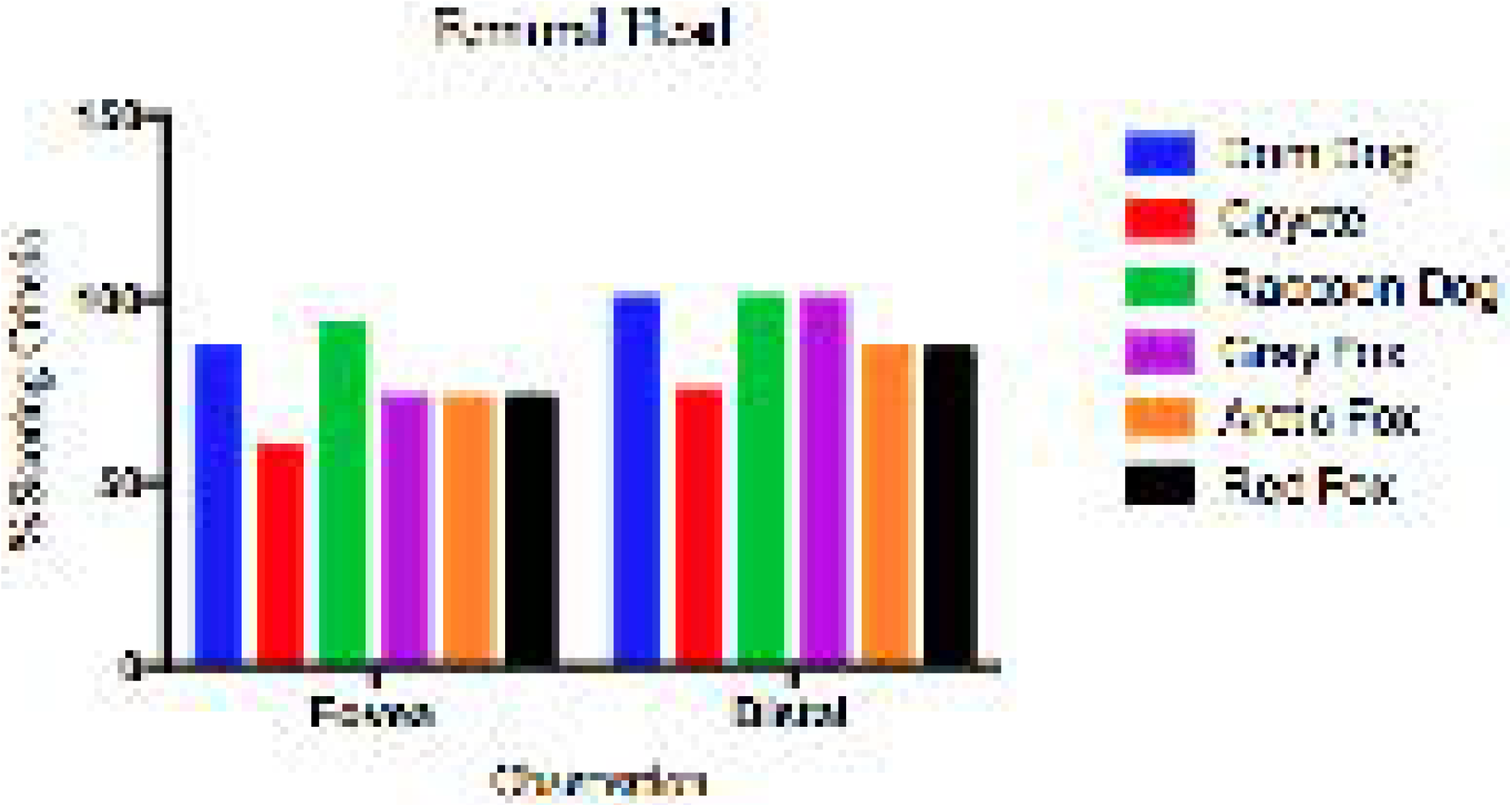

The graphical representations of the outcomes within each of the 5 scoring protocols (fossa, medial articular margin, articular surface, lateral articular margin, and femoral head) reveal some detail.

From Figs. 7 – 11 and Supplement Table 1, the relationships among the taxa can be approximated for each intra-articular structure. Pathology scores for the <u>acetabular fossa</u>, except for uniformly high presence of “accessory fossa osteophytes”, were varied among the taxa (Figs. 1, 7). Pathology scores for “extent” and “character” of the <u>medial articular margin</u> were uniform across taxa, and generally exceeded 75% (Figs. 2, 8). Pathology scores for the <u>acetabular articular surface</u> “character” varied among taxa, but spatial scores were uniform and high for “medial-central-lateral” and for “distribution”. “Cranial-caudal” articular surface spatial scores were high for raccoon dogs, and moderately high for the other taxa (Figs. 3, 9). For the <u>lateral articular margin</u>, pathology scores for “extent” and for “osteophytes” were high, while those for “character” were lower and variable (Figs. 4, 10). Pathology scores for the <u>femoral head</u> varied among taxa for “fovea” and “distal non-articulating surface component” (Figs. 5, 11). Since the raccoon dogs in the database had been farm-maintained and likely were older at harvesting, their appearance of generally higher scores would be expected, where features were not obscured.

## DISCUSSION

Within scoring and statistical limitations, across taxa and across intra-articular structures, at the same loci, the observations revealed high prevalence of significant pathology. From earlier studies, we had observed that pathology at this morphological level largely is incipient-to-subtle (sub-radiographic) (Mustonen et al., 2017; Lawler et al., 2021a, 2021b). However, the more important implication is that all of the taxa clearly have the genomic and/or epigenomic capacity for the responses (defined by its presence). Therefore, a primary question can be phrased: What is (are) the explanation(s) for the high frequency of incipient-to-subtle pathology trait presence across canid taxa, at the same intra-articular loci, whether wild or domesticated?

### General genomic considerations

Graphodatsky et al. (2008) noted that modern canids have “highly rearranged karyotypes”, but nonetheless reveal broad conservation of whole chromosomes. The observation itself suggests complex phylogeny that is centered on gene flow/hybridization. Gene flow is defined as movement of genes among genetically distinguishable populations, resulting in viable offspring (Petit and Excoffier 2009) (Mallet, 2014). Intraspecific gene flow helps to define a species and promote its cohesion. Interspecific gene flow can result in genetic invasion of a species’ genome (introgression), with subsequent effects on species integrity (Petit and Excoffier 2009).

Under certain conditions, gene flow has the potential to enrich variation, increase adaptive phenotypes, and become a stimulus for evolution (Anderson and Stebbins 1954). Such conditions include physical barriers (topography), historical factors (previous colonization, range expansion, glacial isolation), and/or ecological factors (climate, habitat type, prey selection) (Pilot et al. 2006). Another important condition is the transition to a cursorial predation pattern that results in changes in limb skeletal anatomy (Andersson and Werdelin 2003). In turn, divergence of body size can be one influence on prey choice and diet (Seehausen 2013).

More specifically, any analysis of carnivoran appendicular skeletal morphology in terrestrial animals ultimately must relate to locomotor function for survival (successful pursuit and capture of prey). To integrate development, function, and evolution, we present a summary of the phylogenetic history, typical prey species, and locomotor specialization pattern. Carrano (1999) and Martin-Serra et al. (2015) described cursorial (running down prey) versus non-cursorial (pounce and grappling prey with the forelegs) locomotor habit. While these categories are useful conceptually, they should be thought of as representing poles of a continuum.

### Natural history and genomic studies

According to Wang et al. (2008), Family Canidae originated in North America, during the Eocene (c. 40 Mya). Nyakatura and Bininda-Emonds (2019) suggested that, since the initial deep-time divergence (c. 16.5 Mya – 16 Mya) toward the canid lineages was brief in geological time (c. 0.5 Mya), a “genetic blueprint” had to have been established during earlier evolution.

The early diversification yielded three major Subfamilies Hesperocyoninae, Borophaginae, and Caninae (late Oligocene; Wang et al., 2004). The first two were extinct, respectively, by mid-Miocene and early Pleistocene. Genus *Eucyon* (Caninae) canids are thought to have crossed the Beringian land bridge to Asia c. 6 Mya (Wang et al., 2008). Appearances of canids in South America emerged later, during the late Pliocene (Wozencraft 2005, Lucherini and Vidal 2008). The vulpini and canini are thought to have evolved from a split with *Leptocyon* spp. c. 12 Mya – c.11 Mya.

### Tribe Vulpini

Tedford et al. (2009) suggested that the vulpine genera, *Vulpes* and now-extinct *Metalopex*,appeared after c. 9.5 - 9 Mya. Nyakatura and Emonds (2012) suggested a more recent date, after c. 8 Mya. Vulpine phylogeny is confused by the deeply rooted lineages (after c. 16 Mya; Nyakatura and Emonds, 2012) that led to modern *Urocyon* and *Nyctereutes*, with the basal *Urocyon* lineage suggested as the oldest lineage. We suggest that acquiring new paleontological data for deep-time intervening years will be needed to resolve further questions surrounding vulpine phylogeny. For this study, red fox (*Vulpes vulpes*), gray fox (*Urocyon cinereoargenteus*), arctic fox (*Vulpes lagopus*), and raccoon dog (*Nyctereutes procyonoides*) specimens were used because the specific required data were adequate in number.

#### Red fox

Kutschera et al. (2013; mitochondrial DNA) indicated that the red fox was diversifying in Eurasia during the late mid-Pleistocene, with repeated climate-aligned colonizing of some regions. Evidently, red foxes used intermittent land bridges, LGM survival in southern refugia, and natural adaptability, to evolve into an unstructured and panmictic population during the Pleistocene (Kutschera et al., 2013).

Aubry et al. (2009; mitochondrial cytochrome b gene and D-loop) suggested that Holarctic and Nearctic clades diverged c. 0.4 Mya. The Neararctic population has three subclades, one more widespread, dating from c. 45 kya, and two from c. 20 kya. Aubry et al. (2009) added that the Nearctic clades are distinct from the Holarctic clade. Red fox colonizing of North America from Eurasia appears to date from c. 191 – 130 Kya (Illinoian period) (Aubry et al., 2009).

Teacher et al. (2011; mitochondrial cytochrome B and control region DNA) reported no indication of spatial structure, whether in modern or ancient populations. The authors observed no evidence of bottlenecks, no separate lineages or losses of pre-LGM “unique” haplotypes, and low levels of random genetic drift. High long-term dispersal and adaptability occurred with minimal differentiation over wide geography.

Inoue et al. (2007; mitochondrial cytochrome b gene and control region) observed a divergence of c. 1.4 Mya between the Hokkaido and No. Honshu Japanese red fox populations. The larger, more broadly-distributed Group I occurred across Hokkaido and over parts of Eurasia, with Group II on Hokkaido only. Yu et al. (2012; mitochondrial DNA cytochrome b sequences) noted three clades. Clade I is the Hokkaido lineage. Clade II is North American. Clade III includes two lineages with unclear phylogeny. Divergence estimates range between 1.5 Mya and 1.0 Mya.

Red foxes are generalized in terms of morphology and physiology, being slender, long-legged, and relatively large-sized among extant vulpines. They are successful predators and scavengers in many ecological settings (Henry, 1996; Cypher, 2003). Their diet can include small mammals, insects, and fruit, according to availability and necessity (Henry, 1996; Cypher, 2003). The red fox would be considered a low-speed, prolonged trotting type of cursor.

#### Gray fox

The red (*Vulpes lagopus*) and gray (*Urocyon cinereoargenteus*) fox are similar in life habits, although they are not related closely (Henry, 1996; Graphodatsky et al., 2008). They occupy diverse habitats as energetic hunters. The gray fox prefers forest habitats, while the red fox prefers “edge environments” (Henry 1996). The gray fox tree-climbing behavior in forest areas allows access to bird eggs and fruits (Henry 1996). Fritzell et al. (1982) noted that eastern and western (North America) gray foxes prefer forest and scrub ecology, respectively. Based on our observations, gray fox natural behaviors may not contribute additionally to coxofemoral joint pathology, although additional research is advisable because the overall high numbers of pathology observations that we encountered may mask some differences. Martin-Serra (2015) classifies the gray fox as non-cursorial.

Wayne et al. (1997; molecular systematics) stated that the gray fox and the island fox (*U. littoralis*) are the most basal taxa among extant Canidae. Nyakatura and Bininda-Emonds (2012; DNA sequence data) suggest a *U. cinereoargenteus* – U. *littoralis* divergence c. 1.2 Mya. Nyakatura and Bininda-Emonds (2012) also suggest that the lineages leading to *Otocyon* and *Nyctereutes* extend to c. 16 – 15 Mya.

Goddard et al. (2015; mtDNA cytochrome b; coalescent simulations) suggest a deep divergence between eastern and western gray foxes c. 0.5 Mya. Goddard et al. (2015) observed that the modern western *Urocyon cinereoargenteus* could represent a divergence from the earlier *U. galushai* and *U. citrinus*, during the Irvingtonian (1.8 Mya – 0.3 Mya). Nyakatura and Bininda-Emonds (2012) also suggest that lineages leading to *Otocyon* and *Nyctereutes* extend to c. 16 – 15 Mya.

Kumar et al. (2015; tissue transcriptome RNA sequencing) observed lower genetic variability in the arctic fox, compared to the red fox, estimating a divergence c. 2.9 Mya. Vozdova et al. (2016; satellite DNA) noted high overall heterogeneity among red fox, arctic fox, raccoon dog, and domestic dog, although the arctic fox and red fox proved to be related closely. Early in the 20^th^ century, the *American Fur Breeder* (1929) had noted the arctic fox x red fox hybridizing (reported by Gray 1954).

#### Arctic fox

Arctic foxes (*Vulpes lagopus*) have habitat-related natural history limitations and are considered to be low-speed cursors. The rigorous arctic climate dictates shorter life expectancy that may be related to larger litter sizes, between 5 and 15 kits (Henry 1996). Henry (1996) described their primary prey as lemmings and other burrowing mammals, along with forage for seasonal plants, bird eggs, and young of arctic birds. Nearby marine coastal environments provide fish, invertebrates, ocean birds, and some plant material (Henry, 1996; Lawler et al., 2020). According to Henry (1996), summer territories are occupied while food is adequate, such as in coastal habitats. In many other instances, the foxes must travel extensively to find food, and may shadow larger carnivores to scavenge on kills.

Nyakatura and Beninda-Emonds (2012) suggested that the emergence of the arctic fox occurred c. 2.5 Mya, approximately coincident with *Vulpes macrotis*, the kit fox, following several earlier splits among vulpini, although the truly basal vulpine lineages probably date to c. 16 Mya. Tedford et al. (2009), however, place the timing of a vulpine split from canini at c. 11 Mya.

Carmichael et al. (2007; genomic DNA), noted that arctic fox allele frequencies were more homogenous (compared to arctic wolves). No historical bottlenecks were identified, with no presence in Pleistocene refugia. The larger effective population size, high gene flow, longdistance foraging, and lack of population structuring, indicate broad geographic presence of a panmictic taxon.

Mustonen et al. (2017) described skeletal remains of late juvenile farmed arctic foxes, recording incipient pathology matching that observed at intra-articular loci among free-living juvenile or adult arctic foxes (Lawler et al., 2021a; Lawler et al., 2021b). Skeletal remains of free-roaming arctic foxes on St. Paul Is. (Pribilof Is.), Alaska revealed features suggesting that skeletal effects of naturally-occurring malnutrition were superimposed on pre-existing joint pathology (Lawler et al., 2020)

Ploshnitsa et al. (2011) studied MHC genes of two separated arctic fox populations in the Commander Islands. Smaller numbers of MHC genes in one population were associated with decreased fitness, reflected in greater susceptibility to infectious agents and poorer fur quality. The study serves as a good example of fitness losses that can occur, even in subpopulations of panmictic taxa. Population-specific losses in fitness well may be lost to diagnosis, over long geological time. Lotsander et al. (2021) evaluated inbreeding and genetic rescue in a small population of Scandinavian arctic foxes. The authors concluded that genetic rescue via immigration is a complex process with possibly limited population benefit, absent of continuous gene flow.

#### Raccoon dog

Modern raccoon dogs (*Nyctereutes procyonoides*) are rather short-legged, and approximate the size of the smaller foxes (Ward and Wurster-Hill, 1990). According to Nyakatura and Bininda-Emonds (2012), the *Nyctereutes* spp, line emerged early, c. 16 Mya, thus being one of the oldest recognized (and extant) canid genera. This view potentially places these three “long-branch taxa” (authors’ wording), *Nyctereutes* and *Otocyon* as “sister taxa” (authors’ wording) to remaining vulpines, and *Urocyon* as a “sister taxon” to other canids (Nyakatura and Bininda-Emonds, 2012). However, the exact timing and relationships do not seem to be entirely clear at present.

The paleontological record has revealed larger and smaller body forms. It has been suggested that the larger early Pleistocene *N. megamastoiodes* and the mid-Pliocene *N. sinensis* may have been the same animal, with *N. sinensis* evolving to the smaller-sized modern *N. procyonoides* (Kurten 1968).

*Nyctereutes* spp. prefer forested areas around water, with dense undergrowth. Their varied omnivorous diet includes plant, small vertebrates, and invertebrates (Ward and Wurster-Hill, 1990). Novikov (1956) reported significant consumption of fresh water and marine animals in those settings. Thus, as foragers, this species is classified as non-cursorial. By contrast, captive-bred raccoon dogs have been farmed extensively for their pelts (Ward and Wurster-Hill, 1990), and often have been fed to extreme obesity and imbalance of dietary minerals, resulting in obvious nutritional disorders (Lawler et al., 2012).

Considering free-roaming vs. captive-reared vulpines, our coxofemoral joint data reveal few differences (Lawler et al., 2021a; Lawler et al., 2021b). The data support that conserved physiological pathways for growth–development and insult response may overshadow ecological and nutritional effects that are imposed as life events (Tangredi and Lawler, 2019).

### Tribe canini

Nyakatura and Bininda-Emonds (2012; supertree data) suggest a very early group of Canidae (or canid precursors?), c. 16 – 15 Mya. Original known Caninae (*Leptocyon*) were fairly small size and probably did not expand numerically until the approximate time of Borophaginae decline (Tedford et al., 2009, Figs. 1 & 2; paleontological data) from c. 12 Mya. Nyakatura and Bininda-Emonds (2012) suggest the expansion of canini after c. 9.5 Mya. According to Tedford et al. (2009), the earliest tribe canini (*Eucyon*) emerged c. 9 Mya. Nowak (2003) suggested that genus *Canis* emerged over a period of 9 – 4.5 Mya. For our study of canini, coyote (*Canis latrans*) and domestic dog (*C. l. familiaris*) specimens were used.

#### Coyote

Blancan (c. 5 Mya – 2 Mya) coyotes are referred to *C. lepophagus*, and thus the taxon is quite variable (Nowak 2001). *C. lepophagus* or a very close taxon is thought to be the foundation for all modern coyotes. Nowak (2001) suggested that coyotes had developed by the end-Blancan, remaining stable thereafter. Genetic admixture in the wild with gray wolves, red wolves, and domestic dogs is recognized (Nowak 2001).

The highly-adaptable coyote has remained a generalized hunter and an efficient scavenger (Hilton 2001). The less specialized size and body form, throughout its complex history, facilitated an adaptive natural history (Nowak 2001). In particular, Nowak (2001) emphasized smaller average size compared to its canid contemporaries over time, with skull and dental morphology suggesting capacity to use multiple ecologies and food sources. Maintaining these more generalized (and somewhat variable) features thus appears to be a blueprint for long-term success (Nowak 2001).

As a low-speed cursorial, coyote feeding habits include sprinting after running prey, daylight hunting, and quietly stalking small prey (Bekoff 2001). Depending on ecological and geographic circumstance, the coyote diet might vary around deer and livestock kill or carcasses; rabbits, woodchuck, beaver, and muskrat; fruit and insects; and numerous smaller mammals. Coyotes thus take advantage of open or farm land, woodlands, and human-occupied areas (Hilton 2001).

#### Domestic dog

*C. lepophagus* (late Blancan – early Irvingtonian), or a closely related unidentified taxon, appears at or near the significant split within canini that led to *C. latrans* Nowak (2003). The “other arm” of the split probably led to *C. priscolatrans*, with a second significant split during the early Irvingtonian (Nowak 2002, 2003). Nowak (2002) suggested that the second split (early Irvingtonian) likely led to *C. armbrusteri* (late Irvingtonian) and *C. dirus (Aenocyon dirus;* c. 130 kya to 10 kya; Perri et al., 2021). *C. dirus* was the last of the *C. armbrusteri* group, an evolutionary dead end by end-Rancholabrean (Nowak 2003). The gray wolf (and thus the domestic dog) probably descended from a population of Eurasian *C. mosbachensis* or a similar now-extinct taxon that diverged initially around the time of the *C. priscolatrans* lineage split (mid-Irvingtonian) (Nowak 2002, 2003). Nowak (2003) noted that the wolf and coyote diverged during the Pliocene (Late Blancan; c. 2.5 – 1.8 Mya).

The Early Rancholabrean - Holocene Eurasian dominance of *Canis* (Nowak 2003) led eventually to domestication of the gray wolf by humans. Perri et al. (2021; aDNA sequencing) reported that wolves likely were domesticated in Siberia c. 23 kya. This early human-dog relationship would have preceded heretofore-accepted dating for wolf domestication by several thousand years. While retaining much of the wolf ontogeny (Geiger et al. 2016), the modern domestic dog has resulted significantly from selective breeding that resulted in many “modern breeds”, beginning roughly from the Victorian Period of human history. Some modern dog breeds (and those with mixed-breed ancestry) do retain the high-speed cursorial characteristics that were possessed by ancestral populations, but many outcomes of human interventions have been deleterious to dog health and self-sustainability (Lawler 2016).

Lindlbad-Toh et al. (2005) published a dog genome sequence and examined various haplotypes over wide geographic areas. The authors concluded that the data support discovery of specific disease-related genes, although no universal consensus exists on whether the modern domestic dog genome is an optimal model for phylogenetic study.

Finarelli and Goswami (2013) noted that evolutionary influences on specific traits may or may not reflect a given clade’s deeper-time history. The concerns include extinct subclades that may have created admixtures that influenced genomic status, prior to their disappearance. Further, evolutionary trends may be invisible, absent of data from successive intervening taxa (Finarelli and Goswami, 2013). Bergstrom et al. (2020) sequenced 27 ancient Eurasian dog genomes (up to 10.9 kya). All of these were distinct from extant wolves, with gene flow mostly unidirectional as dog to wolf. Thus, the authors suggested that all dogs descend from one wolf population, or population group, that itself became extinct subsequently.

#### Population variability

Further considering genomic aspects, gene flow is movement of genes among genetically distinguishable populations, resulting in viable offspring (Petit and Excoffier 2009) (Mallet, 2014). Viability is an important component in nature, given that an F_1_ generation often can be sterile, due to incompatibility of alleles (Darwin 1876, Mallet 2014). Intraspecific gene flow helps to define a species and promotes its cohesion, whereas interspecific gene flow can result in genetic invasion of a species’ genome (introgression) and affect species integrity (Petit and Excoffier 2009).

The history of Canidae involves long-term gene flow (Gopalakrishnan et al., 2018; Nosil et al., 2021). High gene flow and hybridizing necessarily imply compatible reproductive biology, along with geography (larger range), chronological timing, ancestral admixtures, larger effective population size, long-distance movement, and minimal population structure (Carmichael et al., 2007, Gopalakrishnan et al., 2018). Thus, the “original” blueprint for phenotypes among extant taxa likely extend into deep time, in complicated fashion.

There is a complex interaction among plasticity, selection, and gene flow. For example, high gene flow may tend to maximize fitness to an “average” environmental condition. Also, high plasticity can promote gene flow (Crispo 2008). Thus, hybridization has the potential to enrich variation, increase adaptive phenotypes, and stimulate evolution (Anderson and Stebbins 1954). Under certain conditions, a “hybrid swarm” can result in adaptive radiation of new species (Seehausen 2013, Fontdevila 2019). Such conditions include physical barriers (topography), historical factors (previous colonization, range expansion, glacial isolation), and/or ecological factors (climate, habitat type, prey selection) (Pilot et al. 2006). Divergence of body size can be one influence on prey choice and diet (Seehausen 2013). The transition to a cursorial predation pattern changes prey choice and influences limb skeletal anatomy (Andersson and Werdelin 2003).

### Conclusions

Among all of these taxa, the carnivorous “lifestyle” is rigorous, with the clear exception of the domestic dog. Considering the varied natural histories, it is noteworthy that coxofemoral joint pathology occurred at the same intra- and extra-articular femoral and acetabular loci, whether in juveniles or adults, with only minor differences among taxa. One important practical implication is that joint pathology induced during late juvenile and adult life unavoidably is superimposed on pre-existing but unobserved (sub-radiographic) pathology (Lawler et al., 2021a; Lawler et al., 2021b). This explanation could, in part, account for the often-increasing severity of joint pathology among taxa possessing human-selected traits and their consequences, and for the same observations among wild taxa that survive to advanced age.

## Notes

### Competing Interest Statement

The authors have declared no competing interest.

