## Supplementary figures and images for "Reading the canid skeletal story: Coxofemoral joint pathology, and suggested implications for the phylogenetic and natural history of taxa"

### Supplementary File 1

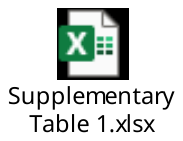
